# ScRNA-seq discover cell cluster change under OAB——ACE2 expression reveal possible alternation of 2019-nCoV infectious pathway

**DOI:** 10.1101/2020.06.05.137380

**Authors:** Boyu Xiang, Xiangyu Hu, Haoxuan Li, Li Ma, Hao Zhou, Ling Wei, Jue Fan, Ji Zheng

## Abstract

**Objective:** Previous study indicated that bladder cells which express ACE2 were a potential infection route of 2019-nCov. This study observed some differences of bladder cell cluster and their ACE2 expression between OAB mice and healthy mice, indicating the change of infectious possibility and pathway under overactive bladder (OAB) circumstance.

**Material and method:** Pubic dataset acquisition was used to get ACE2 expression in normal human bladder and mice bladder (GSE129845). We built up over OAB model and studied the impact on cell typing and ACE2 expression. By way of using single-cell RNA sequencing (scRNA-seq) technique, bladder cell clustering and ACE2 expression in various cell types were measured respectively.

**Result:** In pubic database (healthy human and mice bladder), ACE2 expression in humans and mice is concentrated in bladder epithelial cells. The disappearance of umbrella cells, a component of bladder epithelial, was found in our OAB model. In the two mouse bladder samples, ACE2 expression of epithelial cells is 34.1%, also the highest of all cell types.

**Conclusion:** The disappearance of umbrella cell may alternate the infection pathway of 2019-nCov and relate to the onset and progression of OAB.

## Introduction

In December 2019, a novel coronavirus named 2019-nCoV emerged in Wuhan, China, and swept around the world in just one month. Except remarkable fever and respiratory disorder, urinary system damage was also detected in some 2019-nCoV patients^1, 2^.Though there has been no evidence of positive detection of 2019-nCoV in urine so far, its detection in urine samples of SARS patients hinted the importance of virus-testing in urine samples of 2019-nCoV patients, which showed that the urinary system was a potential route of infection^3, 4^The expression and distribution of receptors determine the pathway of viral infection, which is of great significance for understanding the pathogenesis and designing treatment strategies^5^. 2019-nCov was reported to share the same receptor with SAR-Cov, Angiotensin-converting enzyme 2 (ACE2), and previous study has proven that bladder epithelial cells are potential target cells for a novel coronavirus and the concentration of ACE2 seemed to have a special pattern which shows a decreasing trend from the out layer of the bladder epithelium (umbrella cells) to the inner layer (basal cells) with the intermediate cells in-between^6^.

Studies and a lot of data show that older people with basic diseases have a higher chance of infection and higher mortality if they are infected^7^, so it is very urgent and important to figure out how the underlying disease affects the way to change the 2019-nCov infection pattern. The purpose of our study is to explore cell cluster structure and ACE2 expression pattern in the bladder and use scRNA-seq to infer the effects of human senile lesions, OAB, on umbrella cells and intermediate cells in the bladder epithelium from the results of mice. Since OAB is a common senile disease which leads to expensive medical expenses and can be very authentic to simulate the effects of senile diseases on the bladder epithelium^8^ and it greatly affects the quality of life of the patients, we construct a model by using OAB mice.

Bioinformatics scRNA-seq has been widely used in 2019-nCoV research because it has the ability to analyze gene expression of all cell types in multiple tissues with unbiased high resolution. Because this potential urinary tract infection pathway may be critical to prevent 2019n-Cov, here we used two scRNA-Seq transcriptomes to analyze bladder tissue from normal and OAB mice and combined human and mouse bladder ACE2 expression data from public data base to analyze the impact of underlying disease on this potential infectious system^7^.

## Method

### Pubic dataset acquisition and OAB model construction

Gene expression matrices of scRNA-Seq data from normal humans and mice were downloaded from the Gene Expression Omnibus (GSE129845). We replicated the downstream analysis using the code provided by the author in the original paper. We built up over active bladder model (OAB) and studied the impact on cell typing and ACE2 expression. By way of using scRNA-seq technique, bladder cell clustering and ACE2 expression in various cell types were measured respectively.

### Suspension preparation

Bladder tissue samples were gathered and instantly stored in the GEXSCOPE Tissue Preservation Solution (Singleron Biotechnologies) at 2-8°C. Before dissociating the tissue, the specimens were washed with Hanks Balanced Salt Solution (HBSS) for three times and shred into 1–2 mm pieces. At 37°C in a 15ml centrifuge tube, the tissue pieces were dissolved in 2ml GEXSCOPE Tissue Dissociation Solution (Singleron Biotechnologies) for 15min. After reaction, a 40-micron sterile strainer (Corning) was employed to separate cells from cell debris and other impurities. The cell microspheres were re-suspended in 1ml PBS (HyClone) after centrifugation at 1000 RPM, 4℃, for 5 minutes. 2mL GEXSCOPE Red Blood Cell Lysis Buffer (Singleron Biotechnologies) was added to the cell suspension and incubated in order to remove red blood cells. The mixture was then centrifuged at 1000 rpm for 5 min and the cell pellet resuspended in PBS. The number of cells was calculated automatically by TC20 automatic cell counter (Bio-Rad).

### ScRNA sequencing library preparation

In PBS, the concentration of single-cell suspension was 1×105 cells/ml. According to the manufacture’s references (Singleron GEXSCOPE Single Cell RNA-seq Library Kit, Singleron Biotechnologies), Single cell suspension was loaded onto a microfluidic chip thus building up single cell RNA-seq libraries. The resulting single cell RNA-seq libraries were permuted on an Illumina HiSeq X10 instrument with 150bp paired end reads.

### Primary analysis of scRNA-seq raw sequencing data

By using an internal pipeline, raw reads were processed to generate gene expression matrices. Briefly, we moved out reads without poly-T tails at the intended positions and then extracted each read cell barcode and unique molecular identifier (UMI). Before aligning read two to GRCh38 with ensemble version 92 gene annotations, Adapters and poly-A tails were clipped. Using reads with the same cell barcode, UMI and gene were gathered to generate the number of UMIs per gene per cell. Cell number was then determined based on the inflection point of the number of UMI versus sorted by cell barcode curve.

### Quality control, cell type clustering and visualization

To get access to high quality cells, we discarded cells with less than 200 genes and more than 5000 genes, as well as the ones whose mitochondrial content was higher than 20%. After screening, 7330 cells were kept for the following analysis. The tSNE projection of the data calculation was similar to previously mentioned. Cell clusters with resolution 0.8 were obtained by using Seurat Find clusters function. With cell canonical markers, their types were distributed.

## Results

From dataset (GSE129845), cell types annotated were basically the same as the original research^9^, except neuron of mouse (Figure 1A, 1B, 2A and 2B). The fibroblasts in the red box were annotated as neurons by GPM6A in the original article, but more credible markers (Col3a1, Col1a1, and Dcn) proved that they were fibroblasts. In human bladder, the expression of ACE2 was mainly concentrated in three subtypes of epithelial cells, with a small amount distributed in fibroblasts and monocytes (Figure 1C and D). In mouse bladder, ACE2 also gathered in epithelial cells, but with a higher density than human (Figure 2C and D). As shown in Figure 3, the ACE2 expression in human bladder relatively corresponded to that of mouse bladder.

**Figure 1.**
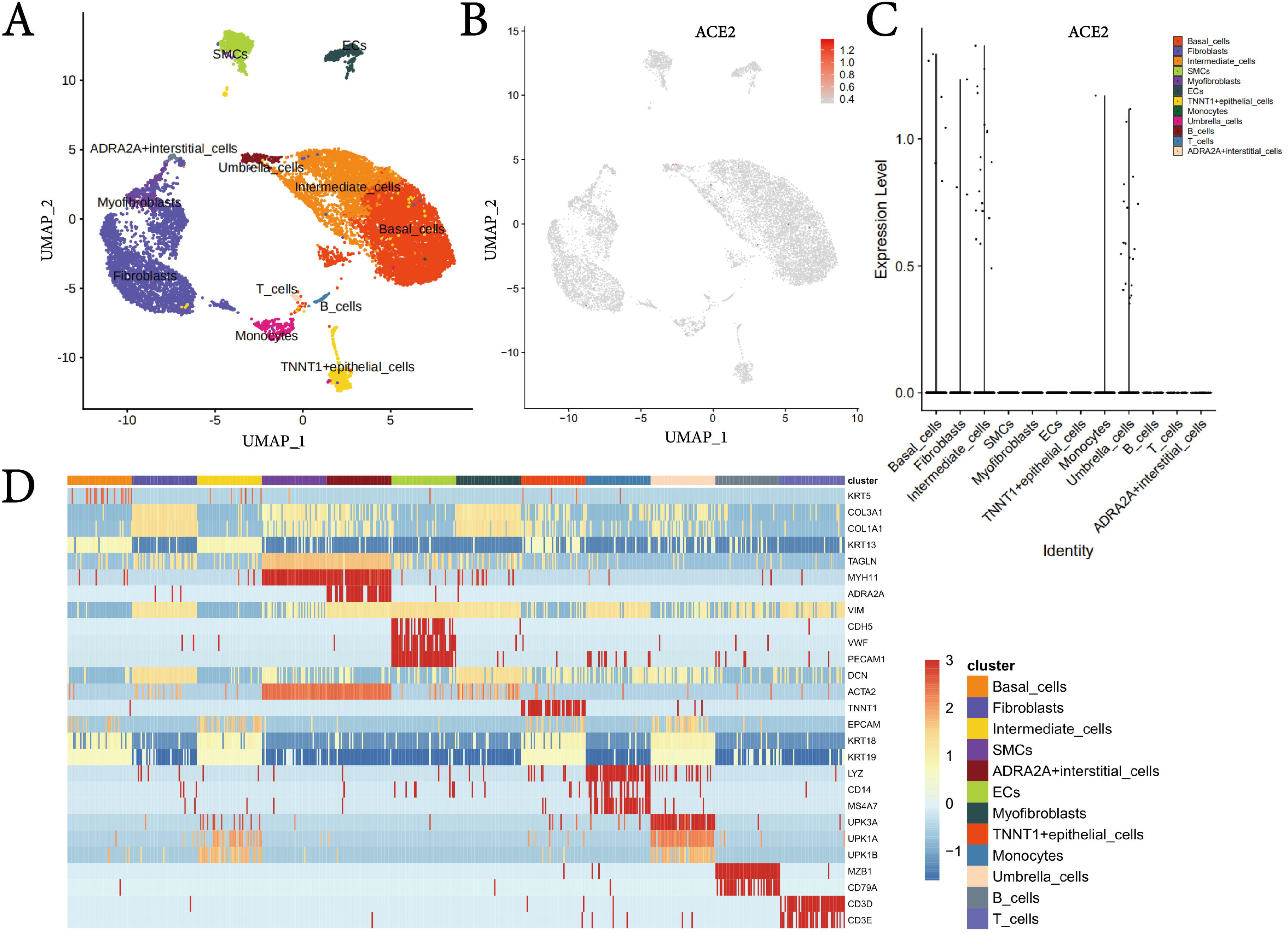
Single cell analysis of the public human bladder data. A) Cell type annotations visualized by UMAP. B) UMAP showing the expression of ACE2 across human bladder. C) Violin plot displaying the ACE2 expression signals of all detected cell types in human bladder. D) Heat map of canonical markers applied for cell type assignment.

**Figure 2.**
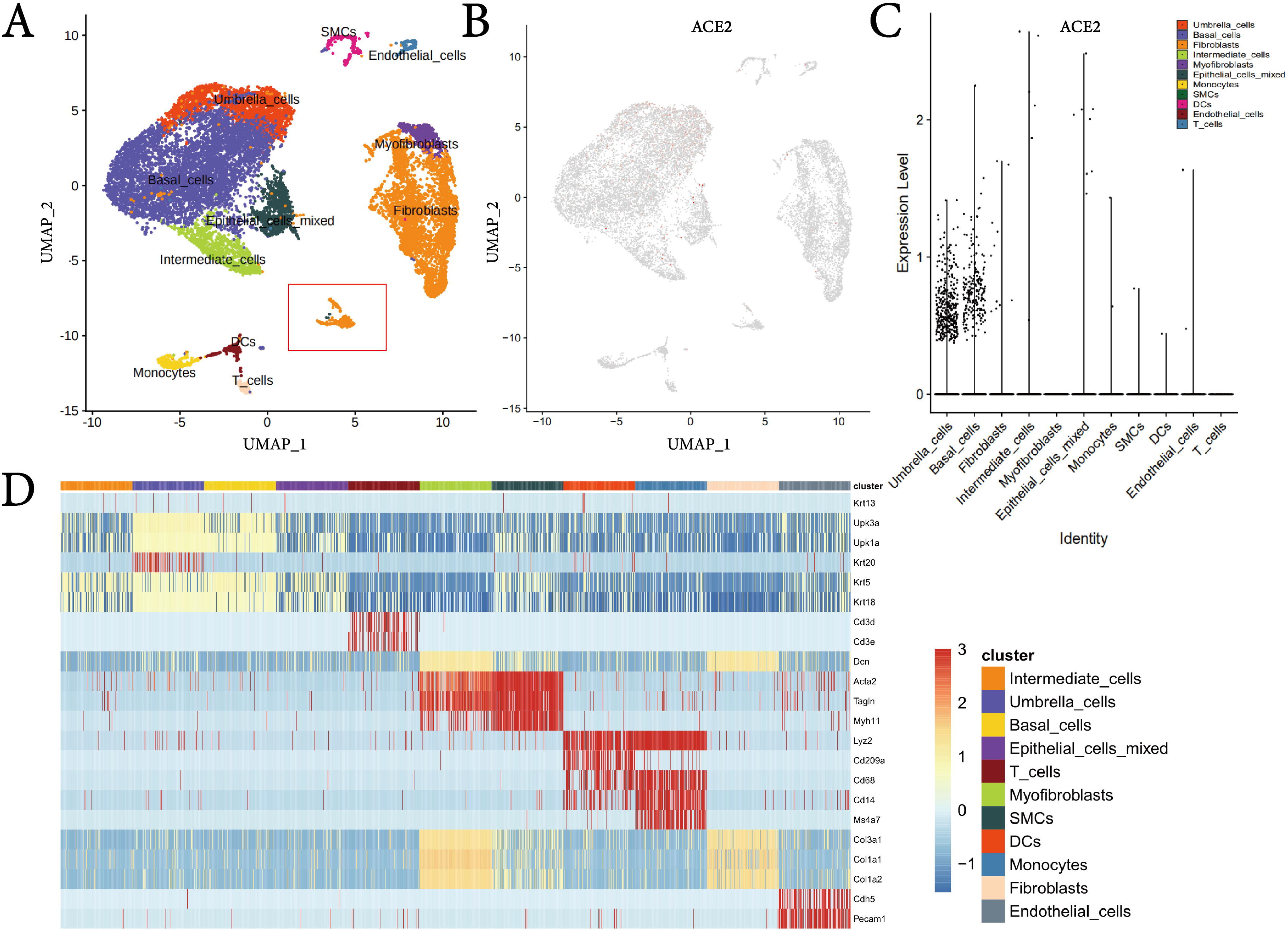
Single cell analysis of the public mouse bladder data. A) Cell type annotations visualized by UMAP. B) UMAP showing the expression of ACE2 across mouse bladder. C) Violin plot displaying the ACE2 expression signals of all detected cell types in mouse bladder. D) Heat map of canonical markers applied for cell type assignment.

**Figure 3.**
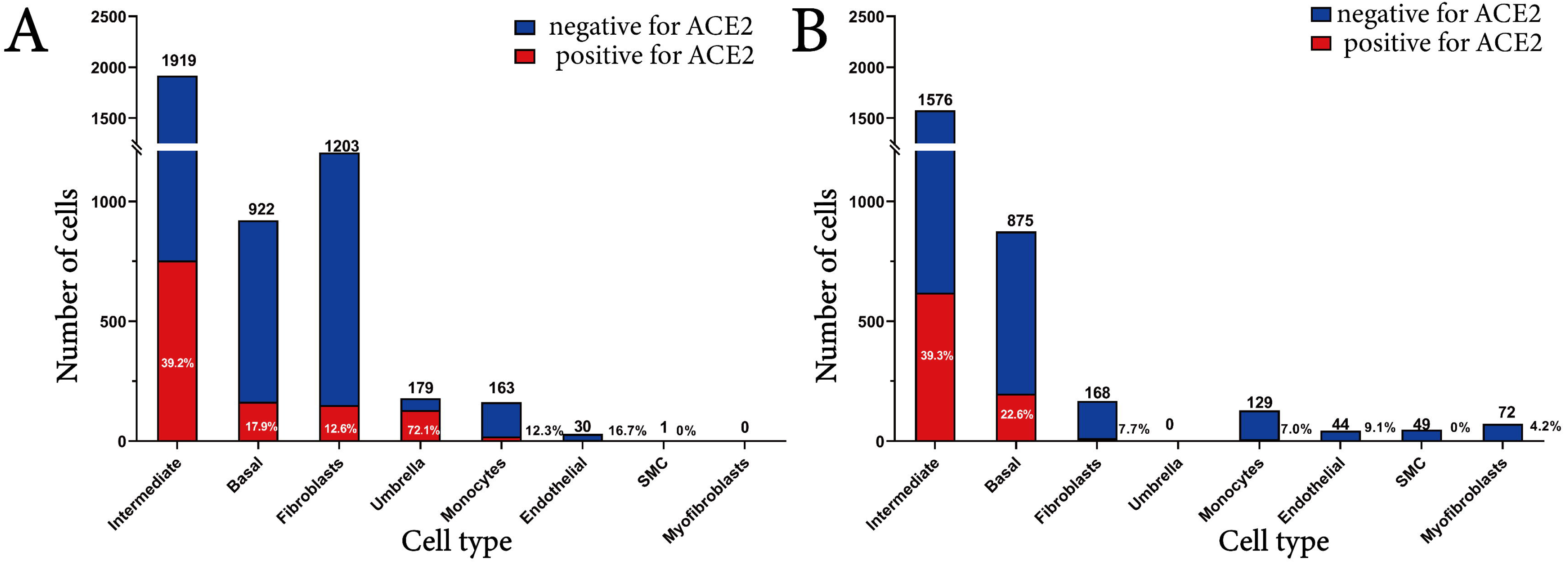
Number of each cell type and percentage of cells expressing ACE2 in healthy human and mouse bladder from public datasets. A) bar chart displaying human bladder cells. B) bar chart displaying mouse bladder cells.

As presented in our independent data (Figure 4), a total of 7330 cells were sequenced, and 8 cell types were distinguished by canonical marker genes. Upk1b, Upk2 and Upk1a were used to identify epithelial cells, the largest part of both samples, along with Krt8 and Krt18 for intermediate cells, Krt20 for umbrella cells. However, the lack of Krt20 signal in OAB sample suggested the disappearance of umbrella cells. Basal cells, which was also a component of epithelium, were separately annotated by Krt5 and Krt17. Fcer1g, Cd14, Cd163, C1qa and Csf1r all together were used to discern monocytes, and Cldn5, Cdh5, Emcn, Plvap and Aqp1 identified Endothelial cells. Fibroblasts, which up-regulated Dcn, Col1a1 and Col1a2, occupied a relatively large part of healthy mouse bladder cells, but decreased significantly in the OAB sample. On the contrary, Myofibroblasts, (Col1a1, Acta2, and Cald1) and Smooth muscle cells (SMC) (Acta2, Tagln, Myh11, and Cnn1) only generated a weak signal in OAB sample. A certain level of ACE2 expression has been detected in almost all cell types of the mouse bladder (Figure 5A-D). As can be seen, most ACE2-expressing cells were intermediate cells and basal cells. However, when it came to the percentage of cells expressing ACE2 (Figure 6), it was umbrella cells that took the dominant position. Fibroblasts, myofibroblasts and SMC were slightly positive for ACE2. The stroma and immune cells, including a small amount of Monocytes and endothelial cells, seemed to down-regulate ACE2 when suffering from OAB.

**Figure 4.**
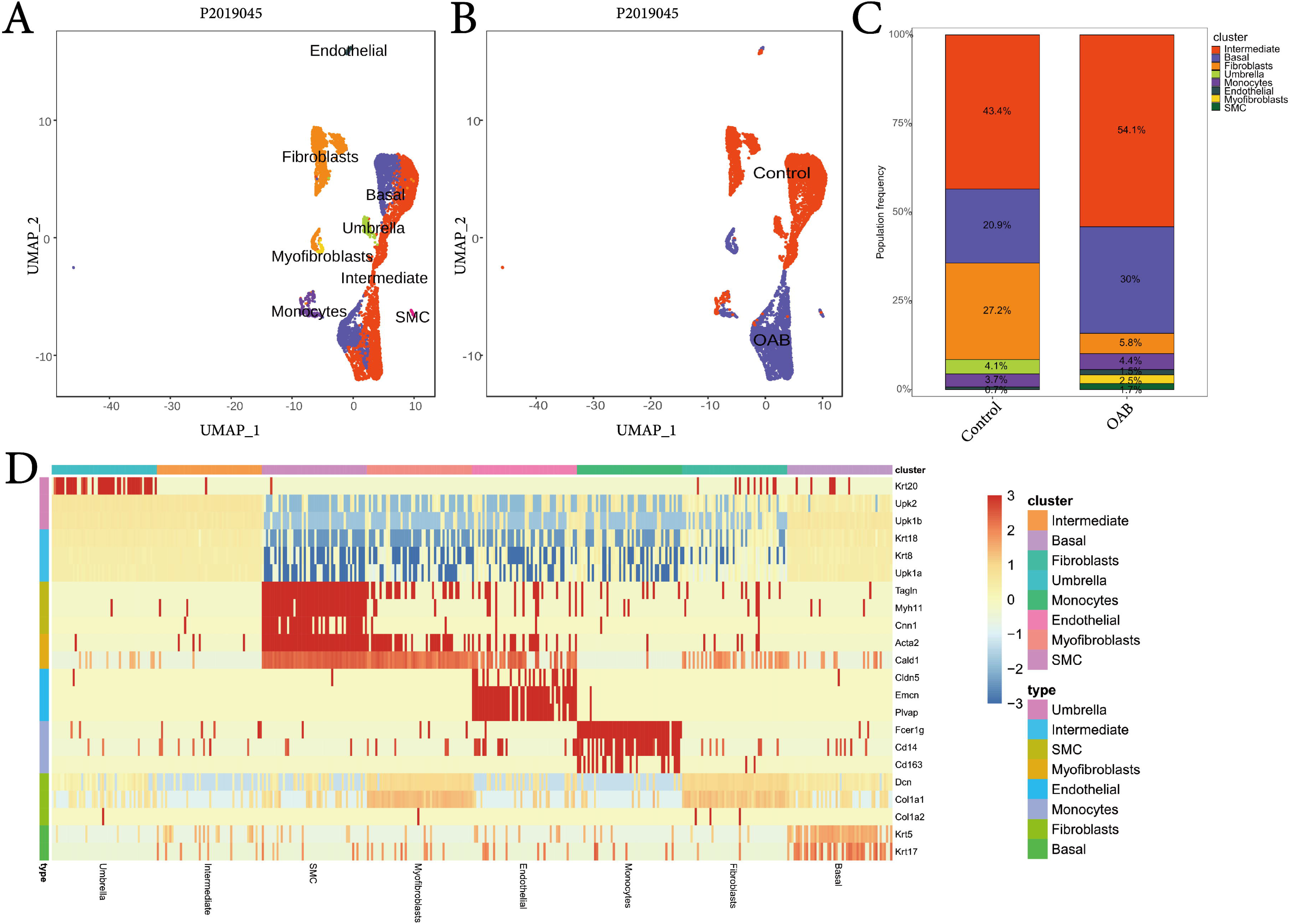
Single cell analysis of the two mouse bladder (control and OAB) samples. A) Cell type annotations of the two samples together visualized by UMAP. B) The sample specific UMAP visualization of two samples. C) The cellular composition and its proportion. D) Heat map of canonical markers applied for cell type assignment.

**Figure 5.**
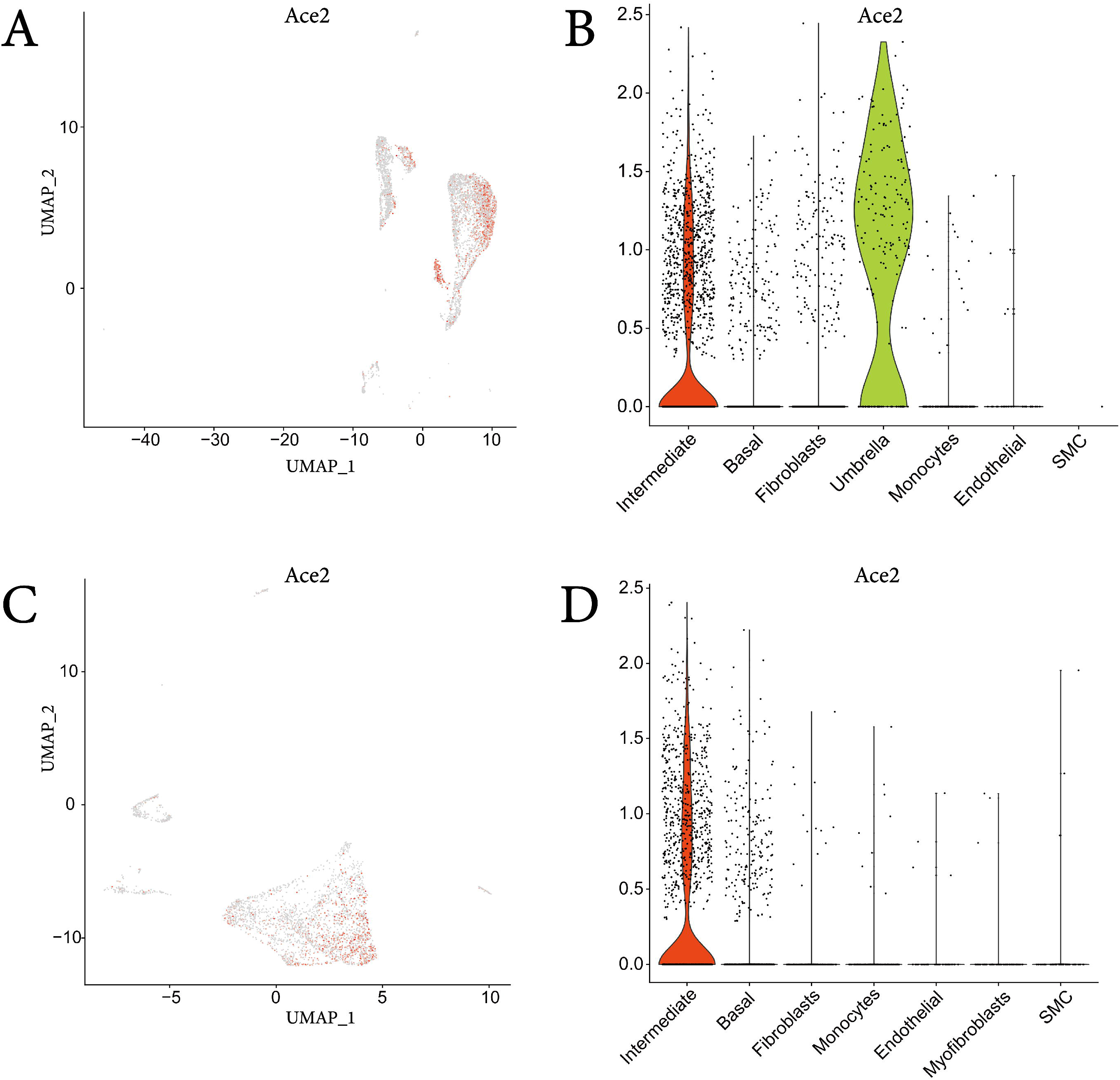
ACE2 expression signals in the two mouse bladder samples. A) UMAP showing the expression of ACE2 across healthy bladder B) Violin plot displaying the ACE2 expression signals of all cell types in control sample. C) UMAP showing the expression of ACE2 across bladder with OAB disease. D) Violin plot displaying the ACE2 expression signals of all cell types in OAB sample.

**Figure 6.**
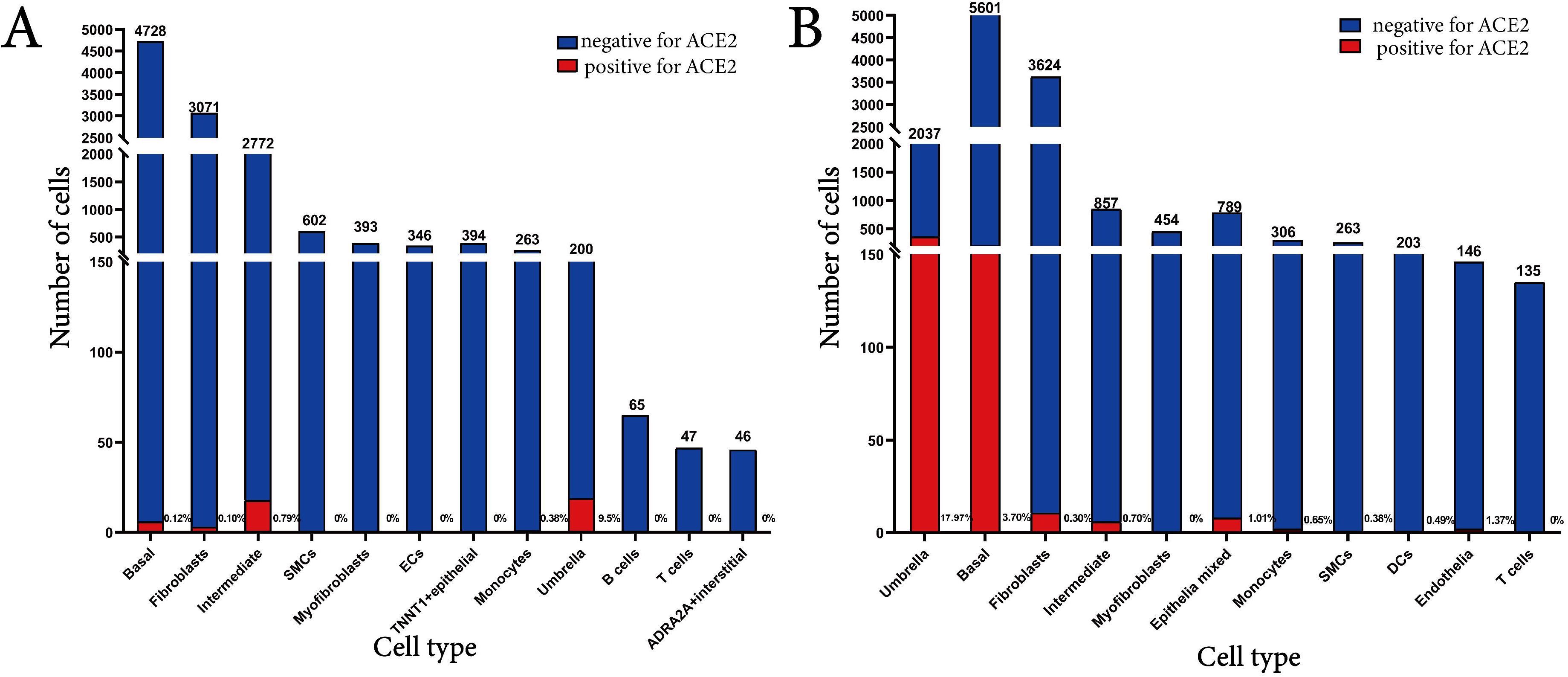
Number of each cell type and percentage of cells expressing ACE2 in healthy mouse and mouse with OAB. A) Bar chart displaying cells of healthy mouse. B) Bar chart displaying cells of mouse with OAB.

## Discussion

According to the latest scRNA-seq studies, ACE2 proved detectable in respiratory, digestive and urinary systems^6, 10, 11^, indicating these systems were potential targets for 2019-nCoV dissemination. Lin, et al have revealed that in bladder, the expression level of ACE2 was relatively high in epithelial cells, especially umbrella cells^6^, which indicated umbrella cells or epithelial cells were more likely to be infected by 2019-nCoV. As we know, the bladder epithelium is a transitional epithelium with high degeneration and rapid cell regeneration, thus bladder diseases like OAB may have an impact on it. In our study, we sought to figure out the changes of scRNA-Seq transcriptome under OAB circumstance. However, healthy or non-cancerous human bladder samples are usually unobtainable. We compared each cell type across species, we found that most human and mouse bladder cells were relatively well correlated. By comparing ACE2 expression profiles, we also found similarities: The highest ACE2 concentration existed in epithelial cells of both species. As an alternative, Mice model was reliable to evaluate the effects of OAB on cell clustering and 2019-nCoV infection risk.

We identified the cells to construct the cell map, and then further clustered the fibroblasts and epithelial cells. We firstly discovered that OAB influenced the subtypes of fibroblasts: Significantly increasing the proportion of SMC. More surprisingly, umbrella cells, which should exist in the superficial layer of epithelial tissue, disappeared (or decreased to under the detectable level) in OAB sample, while the ratio of the rest two cell types (intermediate cells : basal cells) remained rather close (2.08 of control, 1.80 of OAB). It seems that umbrella cells were eliminated, but other epithelial components remain intact.

Umbrella cells are small in quantity but functionally significant. As known, bladder epithelial cells are continuously exposed to mechanical forces. Previous research has revealed that bladder stretch modulated the cytomembrane surface area by coordinating exocytosis and endocytosis at the apical membrane of umbrella cells^12^. Despite the tight barrier function of the apical membrane, it was still highly hypercompliant^13^. To adapt to membrane surface area change, the apical junctional ring (AJR) which lay in the luminal surface (umbrella cells) of the bladder, expanded during bladder filling and contract upon excreting^14^. These closelyconnection between umbrella cells and bladder activities indicated that the disappearance of umbrella cells is likely to be the pathogenesis or pathological symptoms of OAB. In addition, as the bladder epithelium was a physiological barrier that protected bladder against infection, toxins and material exchange, disappearance of the first layer (umbrellas cells) may cause susceptibility to various pathogens. Considering that OAB was a common disease in the aged, the elderly may need to pay more attention to avoiding urinary tract infections.

Based on single cell analysis, ACE2 expression patterns in the two mouse bladders were summarized. As mentioned before, umbrella cells disappeared under OAB circumstance. Coincidentally, umbrella cells were the most-highly-expressed cell type in the bladder, with a much higher expression percentage (72.07%) than the other cell types, followed by its neighbors (39.28% of intermediate cells, 20.20% of basal cells).The situation of the myofibroblasts is just opposite to the umbrella cells. Along with the cell number increase under OAB circumstance, the ACE2 expression distribution in the myofibroblasts added. Some slight up-regulation or down-regulation of ACE2 was also observed. We do not yet know how these changes relate to OAB, but OAB may alter the infection pathway under these expression pattern changes. Such alterations may not only lead to different susceptibility and disease severity, but also influence the therapeutic effect of general treatment regimens.

However, analysis of different datasets showed that ACE2 expression in bladder was not so stable. Taking umbrella cells for example, for human, the percentage of cells expressing ACE2 was 1.3% in GSE108097^6^ and 9.5% in GSE129845; for healthy mice, it was 18.0 in GSE129845 and 72.1% in our independent unpublished data. Different analytical methods or individual differences may cause these distinct differences. To acquire more precise data, mice batch should be unified.

2019 novel coronavirus pneumonia and severe acute respiratory syndrome (SARS) were both characterized by respiratory complications. However, other organ dysfunctions reflected that 2019-nCoV and SARS-CoV were able to spread elsewhere, including urinary system^15, 16^. SARS-CoV was detectable in the urine samples^17^. More notably, even in convalescent patients’ urine, cultivable SARS-CoV could be detected for longer than 4 weeks^18^, which means their urine might be risky of infection for a lasting time. Our researches also provided single-cell evidence for potential 2019-nCoVinfection of the urinary tract. Recently, 2019-nCoV has been spreading rapidly in countries outside China. The protection of the respiratory tract is certain to be taken seriously, while the protection of the urinary tract is easily overlooked. We suggest patients’ urine be safely disposed and public toilets in severe epidemic areas be regularly disinfected or closed. Still, it remains obligatory to detect 2019-nCoV in the urine specimens.

Yet for all that promise, our method is far from being the whole answers. The results from mouse scRNA-seq were all side evidence without direct relation with clinical features of 2019-nCoV. In addition, we only detected a limited number of mouse bladder samples, rather than hACE2 transgenic mice or human samples. Li et al has reported that compared to human’s ACE2, mice’s ACE2 is less efficient in SARS-CoV binding. Due to differences in molecular structure, such a binding efficiency difference may also appear in 2019-nCoV. Beyond that, the pathogenicity of 2019-nCoV was only clarified in hACE2 mice^19^, so normal mouse model may fail to simulate pathogenesis and therapeutics of 2019 novel coronavirus pneumonia. Further studies should turn to hACE2 mice or humans and increase the sample size.

## ACKNOWLEDGEMENTS

This work was supported by Research Project of Chinese Society of Academic Degrees And Graduate Education (Grant No. 2015Y0405); Key Project of Graduate Education and Teaching Reform of Army Medical University (Grant No. 2018yjgA006)

## Reference

1. H Nishiura, S Jung, NM Linton, et al. The Extent of Transmission of Novel Coronavirus in Wuhan, China, 2020. Journal of clinical medicine 2020;9(2):330. doi:10.3390/jcm9020330

2. Q Li, X Guan, P Wu, et al. Early Transmission Dynamics in Wuhan, China, of Novel Coronavirus-Infected Pneumonia. The New England journal of medicine 2020;382:1199–1207. doi: 10.1056/NEJMoa2001316

3. SKP Lau, XY Che, PCY Woo, et al. SARS coronavirus detection methods. Emerging Infectious Disease 2005;11(7):1108–1111. doi: 10.3201/eid1107.041045

4. CM Chu, WS Leung, VCC Cheng, et al. Duration of RT-PCR positivity in severe acute respiratory syndrome. European Respiratory Journal 2005;25(1):12–14. doi: 10.1183/09031936.04.00057804

5. Y Zhao, Z Zhao, Y Wang, et al. Single-cell RNA expression profiling of ACE2, the putative receptor of Wuhan 2019-nCov. BioRxiv 2020. doi: 10.1101/2020.01.26.919985

6. W Lin, L Hu, Y Zhang, et al. Single-cell Analysis of ACE2 Expression in Human Kidneys and Bladders Reveals a Potential Route of 2019-nCoV Infection. bioRxiv 2020. doi: 10.1101/2020.02.08.939892

7. Emerging understandings of 2019-nCoV. Lancet 2020;395(10221):311. doi: 10.1016/S0140-6736(20)30186-0

8. J Zheng, H Zhou, M Yang et al. Reduced Ca2+ spark activity contributes to detrusor overactivity of rats with partial bladder outlet obstruction. Aging 2020;12(5):4163–4177. doi: 10.18632/aging.102855

9. Z Yu, J Liao, Y Chen et al. Single-cell transcriptomic map of the human and mouse bladders. Journal of the American Society of Nephrology 2019. 30 (11) 2159–2176. doi: 10.1681/ASN.2019040335

10. Y Zhao, Z Zhao, Y Wang, et al. Single-cell RNA expression profiling of ACE2, the putative receptor of Wuhan 2019-nCov. bioRxiv 2020. doi: 10.1101/2020.01.26.919985

11. H Zhang, Z Kang, H Gong, et al. The digestive system is a potential route of 2019-nCov infection: a bioinformatics analysis based on single-cell transcriptomes. BioRxiv 2020. doi: 10.1101/2020.01.30.927806

12. ST Truschel, E Wang, WG, et al. RuizStretch-regulated Exocytosis/Endocytosis in Bladder Umbrella Cells. Molecular Biology of the Cell2002;13(3):830–846. doi: 10.1091/mbc.01-09-0435

13. JC Mathai, EH Zhou, W Yu, et al. Hypercompliant Apical Membranes of Bladder Umbrella Cells. Biophysical journal 2014;107(6):1273–1279. doi: 10.1016/j.bpj.2014.07.047

14. AF Eaton, DR Clayton, WG Ruiz, et al. Expansion and contraction of the umbrella cell apical junctional ring in response to bladder filling and voiding. Molecular Biology of the Cell; 2019;30(16):2037–2052. doi: 10.1091/mbc.E19-02-0115

15. Z Li, M Wu, J GuoCaution, et al. Cautionon on Kidney Dysfunctions of 2019-nCoV Patients. MedRxiv, 2020. doi: 10.1101/2020.02.08.20021212

16. KH Chu, WK Tsang, CS Tang, et al. Acute renal impairment in coronavirus-associated severe acute respiratory syndrome. Kidney international 2005;67(2):698–705. doi: 10.1111/j.1523-1755.2005.67130.x

17. SKP Lau, XY Che, PCY Woo, et al. SARS Coronavirus Detection Methods. Emerging Infectious Diseases 2005;11(7):1108–1111. doi: 10.3201/eid1107.041045

18. D Xu, Z Zhang, L Jin, et al. Persistent shedding of viable SARS-CoV in urine and stool of SARS patients during the convalescent phase. European Journal of Clinical Microbiology and Infectious Diseases 2005;24:165–171. doi: 10.1007/s10096-005-1299-5

19. L Bao, W Deng, B Huang, et al. The Pathogenicity of SARS-CoV-2 in hACE2 Transgenic Mice. bioRxiv 2020. doi: 10.1101/2020.02.07.939389

